# Advanced diffusion imaging in grey matter reflects individual differences in cognitive ability in older adults

**DOI:** 10.1101/2025.05.26.656181

**Authors:** Adam Kimbler, Craig EL Stark

## Abstract

Diffusion Weighted Imaging is a tool that can non-invasively provide insights into the microstructure of a given brain region. Various advanced techniques exist within the diffusion weighted imaging space that each provide valuable insights into different aspects of microstructure. In the following study, we sought to examine whether the combination of derived diffusion metrics (tensors, neurite orientation dispersion and density imaging (NODDI), and mean apparent propagator (MAP) MRI) in grey-matter regions could reliably predict cognitive performance in older adults, and whether these findings were replicable across datasets. First, we demonstrated that all combinations of diffusion metrics could reliably determine participant characteristics and were significant predictors of age. Second, we found that a combination of Tensor, NODDI, and MAP-MRI metrics within the hippocampus could predict RAVLT performance in older adults above and beyond any combination of two metrics alone. We also found these diffusion metrics were able to reliably predict RAVLT performance, but not Trails B or Digit Symbol Substitution Task performance. We also found that these same combinations of metrics could predict working memory performance, but not memory performance within a region associated with working memory (Brodmann Areas 9 and 46). Taken together, these findings indicate that these diffusion metrics provide valuable information on grey-matter microstructure independent of one another, and that the ability to obtain both NODDI and MAP-MRI based information from multi-shell diffusion scans more than justifies the added length.

## 1 Introduction

Magnetic Resonance Imaging (MRI) is an invaluable tool for detecting differences in the brain that underly behavioral, cognition, and pathology. Diffusion Weighted Imaging, a subcategory of MRI that focuses on the diffusion of water molecules across the brain (Stejskal & Tanner, 1965), has frequently been used as a tool to examine the underlying microstructure of the brain, often highlighting the role of white matter tracts in brain function. Diffusion tensor metrics and tractography make the assessment of axonal integrity connectivity between regions relatively straightforward (Assaf & Pasternak, 2008; Sasson et al., 2010; Thomason & Thompson, 2011; Madden et al., 2012). The detection of such minute changes in structure is crucial for the understanding and treatment of different types of pathology, such as Alzheimer’s (Weston et al., 2015) and Parkinson’s (Y. Zhang & Burock, 2020). While most studies tend to focus on changes in white-matter structure using diffusion MRI, research into grey matter microstructure to predict cognition has made great strides in recent years (Budde & Annese, 2013; Aggarwal et al., 2015; Colgan et al., 2016; Assaf, 2019; Radhakrishnan et al., 2020; Venkatesh et al., 2020; Radhakrishnan et al., 2022)

More advanced techniques such as neurite orientation and dispersion and density imaging (NODDI: H. Zhang et al., 2012) can help elucidate the different microstructural components of cells within a given voxel, providing us the estimates of the extracellular and intracellular components and the free water diffusing within the voxel. A more recent tool within diffusion MRI is Mean Apparent Propagator (MAP) MRI (Özarslan et al., 2013), which represents 3D q-space via an analytical series expansion, allowing for use to understand both Gaussian and non-Gaussian diffusion in complex tissue microstructures, allowing previously unanalyzed microstructures to be mapped. While previous work from our lab and others has shown that NODDI metrics in grey matter can be used to predict cognitive performance above and beyond tensors and volume alone (Kamiya et al., 2020; Radhakrishnan et al., 2022), and other work has demonstrated that regional MAP-MRI metrics can predict memory performance on their own (Singh et al., 2024), no study has examined whether combinations of these tensor, NODDI, and MAP-MRI metrics provide an increase in prediction of memory performance.

In the current study, we sought to examine whether diffusion derived metrics (tensors, NODDI, MAPMRI) in grey matter can be used to predict cognitive performance in older adults. We sought to replicate previous findings from our lab indicating that hippocampal diffusion metrics can predict age in both our previously used dataset and an additional dataset (ADNI). Further, we sought to replicate our findings that hippocampal diffusion can predict memory performance on a memory task across multiple datasets, and whether the addition of MAP-MRI based metrics can provide additional predictive utility above and beyond tensors and NODDI alone. Finally, we sought to extend this by assessing whether there is a structure-function relationship present such that hippocampal diffusion metrics would be specifically related to memory tasks and whether other regions’ diffusion metrics might predict individual differences in other cognitive domains.

## 2 Methods

### 2.1 Dataset 1

#### 2.1.1 Participants

Participants in Dataset 1 consisted of 71 older adults recruited from a study at the University of California, Riverside (Radhakrishnan et al., 2022). We then filtered out subject IDs that had a Rey Auditory Verbal Learning delay score of less than 5 (n=15). The final sample consisted of 56 older adults (73.51±6.31 years, 42 female). All participants provided informed consent before participation in this study and were compensated for their time. All experimental procedures were approved by the University of California, Riverside Review Board.

#### 2.1.2 MR Acquisition

Data was acquired on Siemens Prisma 3T MRI scanner (Siemens Healthineers, Malvern, PA with a 32-channel receive-only head coil. Head movements were minimized with fitted padding.

##### T1w

Structural images were acquired using a magnetization-prepared rapid gradient echo (MP-RAGE) protocol (echo time (TE)/repetition time (TR) = 2.72/2400 ms, 208 axial slices, FOV = 300 x 320 mm, GRAPPA acceleration factor = 2, and in 0.8 mm isotropic resolution) that resulted in a T1-weighted image.

##### DWI

Two multi-shell (6 0 s/mm^2^, 64 1500 s/mm^2^, 64 3000 s/mm^2^ shells) sequences were acquired with the following diffusion parameters: TE/TR = 102/3500 ms, FOV = 212 × 182 mm, 64 axial slices, Acquisition time = 10 min and 57 s, multi-band acceleration factor = 4 and in 1.7 mm isotropic resolution. Each acquisition was acquired on the axial plane, with the first acquisition being in the anterior-posterior direction and the second in the posterior-anterior direction.

#### 2.1.3 Anatomical data preprocessing

The T1-weighted (T1w) image was corrected for intensity non-uniformity (INU) using N4BiasFieldCorrection (Tustison et al., 2010, ANTs 2.3.1), and used as T1w-reference throughout the workflow. The T1w-reference was then skull-stripped using antsBrainExtraction.sh (ANTs 2.3.1), using OASIS as target template. Spatial normalization to the ICBM 152 Nonlinear Asymmetrical template version 2009c (Fonov et al., 2009, RRID:SCR_008796) was performed through nonlinear registration with antsRegistration (ANTs 2.3.1, RRID:SCR_004757, Avants et al., 2008), using brain-extracted versions of both T1w volume and template. Brain tissue segmentation of cerebrospinal fluid (CSF), white-matter (WM) and gray-matter (GM) was performed on the brain-extracted T1w using FAST (FSL 6.0.3:b862cdd5, RRID:SCR_002823, Y. Zhang et al., 2001).

#### 2.1.4 Diffusion data preprocessing

Images were grouped into two phases encoding polarity groups. Any images with a b-value less than 100 s/mm^2 were treated as a *b*=0 image. MP-PCA denoising as implemented in MRtrix3’s dwidenoise (Veraart et al., 2016) was applied with a 5-voxel window. After MP-PCA, Gibbs unringing was performed using MRtrix3’s mrdegibbs (Kellner et al., 2016). Following unringing, B1 field inhomogeneity was corrected using dwibiascorrect from MRtrix3 with the N4 algorithm (Tustison et al., 2010). After B1 bias correction, the mean intensity of the DWI series was adjusted so all the mean intensity of the b=0 images matched across each separate DWI scanning sequence. Both distortion groups were then merged into a single file, as required for the FSL workflows. After B1 bias correction, the mean intensity of the DWI series was adjusted so all the mean intensity of the b=0 images matched across each separate DWI scanning sequence. Both distortion groups were then merged into a single file, as required for the FSL workflows.

FSL (version 6.0.3:b862cdd5)’s eddy was used for head motion correction and Eddy current correction (Andersson & Sotiropoulos, 2016). Eddy was configured with a *q*-space smoothing factor of 10, a total of 5 iterations, and 1000 voxels used to estimate hyperparameters. A linear first level model and a linear second level model were used to characterize Eddy current-related spatial distortion. *q*-space coordinates were forcefully assigned to shells. Field offset was attempted to be separated from subject movement. Shells were aligned post-eddy. Eddy’s outlier replacement was run (Andersson et al., 2016). Data were grouped by slice, only including values from slices determined to contain at least 250 intracerebral voxels. Groups deviating by more than 4 standard deviations from the prediction had their data replaced with imputed values. Data was collected with reversed phase-encode blips, resulting in pairs of images with distortions going in opposite directions. Here, multiple DWI series were acquired with opposite phase encoding directions, so b=0 images were extracted from each. From these pairs the susceptibility-induced off-resonance field was estimated using a method similar to that described in (Andersson et al., 2003). The fieldmaps were ultimately incorporated into the Eddy current and head motion correction interpolation. Final interpolation was performed using the Jacobian modulation method to account for signal pile-up/dilution caused by local stretching/compression (Andersson & Sotiropoulos, 2015).

Several confounding time-series were calculated based on the preprocessed DWI: framewise displacement (FD) using the implementation in *Nipype* (following the definitions by (Power et al., 2014). The head-motion estimates calculated in the correction step were also placed within the corresponding confounds file. Slice-wise cross correlation was also calculated. The DWI time-series were resampled to ACPC, generating a *preprocessed DWI run in ACPC space* with 2mm isotropic voxels.

Many internal operations of *QSIPrep* use *Nilearn* 0.8.1 (Abraham et al., 2014, RRID:SCR_001362) and *Dipy* (Garyfallidis et al., 2014). For more details of the pipeline, see the section corresponding to workflows in *QSIPrep*’s documentation.

### 2.2 Dataset 2

#### 2.2.1 Participants

Data used in the preparation of this article were obtained from the Alzheimer’s Disease Neuroimaging Initiative (ADNI) database (adni.loni.usc.edu). ADNI was launched in 2003 as a public-private partnership, led by Principal Investigator Michael W. Weiner, MD. The primary goal of ADNI has been to test whether serial magnetic resonance imaging (MRI), positron emission tomography (PET), other biological markers, and clinical and neuropsychological assessment can be combined to measure the progression of mild cognitive impairment (MCI) and early Alzheimer’s disease (AD). Participants in Dataset 2 consisted of 223 older adults from the Alzheimer’s Disease Neuroimaging Initiative 3 (ADNI3) dataset that had an available multi-shell diffusion MRI acquisition obtained on a Siemens scanner. The data were accessed and downloaded on January 22^nd^, 2024. After obtaining this initial dataset, we filtered out subjects that did not have either the RAVLT or Digit Symbol Substitution Task information available (n=26). We then filtered out all subject IDs that had a diagnosis other than Cognitively Normal (n=86) to keep the data in line with that acquired in Dataset 1. Finally, to further keep the datasets similar, we then filtered out subject IDs that had a Rey Auditory Verbal Learning delay score of less than 5 (n=32). Additionally, one subject was excluded due to low quality segmentation. This resulted in a final sample of 78 subjects (Age=73.13±8.22, 52 female).

#### 2.2.2 MR Acquisition

##### T1w

Structural images were acquired using a magnetization-prepared rapid gradient echo (MP-RAGE) protocol (echo time (TE)/repetition time (TR) = 2.98/2300 ms, 208 axial slices, FOV = 240 x 256 mm, GRAPPA acceleration factor = 2, and in 1.0 mm isotropic resolution) that resulted in a T1-weighted image.

##### DWI

One multi-shell (13 0 s/mm^2^, 6 500 s/mm^2^, 48 1000 s/mm^2^, and 60 2000 s/mm^2^ shells) sequence was available with the following diffusion parameters: TE/TR = 71/3400 ms, FOV = 116 × 116 mm, 81 axial slices, acquisition time = 10 min and 57 s, multi-band acceleration factor = 3 and in 2.0 mm isotropic resolution. The acquisition was acquired on the axial plane with a phase-encoding direction of posterior-anterior.

#### 2.2.3 *Anatomical* data preprocessing

The T1-weighted (T1w) image was corrected for intensity non-uniformity (INU) using N4BiasFieldCorrection (Tustison et al., 2010, ANTs 2.3.1), and used as T1w-reference throughout the workflow. The T1w-reference was then skull-stripped using antsBrainExtraction.sh (ANTs 2.3.1), using OASIS as target template. Brain surfaces were reconstructed using recon-all (FreeSurfer 6.0.1, RRID:SCR_001847, (Dale et al., 1999), and the brain mask estimated previously was refined with a custom variation of the method to reconcile ANTs-derived and FreeSurfer-derived segmentations of the cortical gray-matter of Mindboggle (RRID:SCR_002438, (Klein et al., 2009)). Spatial normalization to the ICBM 152 Nonlinear Asymmetrical template version 2009c (Fonov et al., 2009, RRID:SCR_008796) was performed through nonlinear registration with antsRegistration (ANTs 2.3.1, RRID:SCR_004757, Avants et al., 2008), using brain-extracted versions of both T1w volume and template. Brain tissue segmentation of cerebrospinal fluid (CSF), white-matter (WM) and gray-matter (GM) was performed on the brain-extracted T1w using FAST (FSL 6.0.3:b862cdd5, RRID:SCR_002823, Y. Zhang et al., 2001).

#### 2.2.4 Diffusion data preprocessing

Diffusion processing was done using *QSIPrep.* Many internal operations of *QSIPrep* use *Nilearn* 0.8.1 (Abraham et al., 2014, RRID:SCR_001362) and *Dipy* (Garyfallidis et al., 2014). For more details of the pipeline, see the section corresponding to workflows in *QSIPrep*’s documentation.

Any images with a b-value less than 100 s/mm^2 were treated as a *b*=0 image. MP-PCA denoising as implemented in MRtrix3’s dwidenoise (Veraart et al., 2016) was applied with a 5-voxel window. After MP-PCA, Gibbs unringing was performed using MRtrix3’s mrdegibbs (Kellner et al., 2016). Following unringing, B1 field inhomogeneity was corrected using dwibiascorrect from MRtrix3 with the N4 algorithm (Tustison et al., 2010). After B1 bias correction, the mean intensity of the DWI series was adjusted so all the mean intensity of the b=0 images matched across each separate DWI scanning sequence.

FSL (version 6.0.3:b862cdd5)’s eddy was used for head motion correction and Eddy current correction (Andersson & Sotiropoulos, 2015). Eddy was configured with a *q*-space smoothing factor of 10, a total of 5 iterations, and 1000 voxels used to estimate hyperparameters. A linear first level model and a linear second level model were used to characterize Eddy current-related spatial distortion. *q*-space coordinates were forcefully assigned to shells. Field offset was attempted to be separated from subject movement. Shells were aligned post-eddy. Eddy’s outlier replacement was run (Andersson et al., 2016). Data were grouped by slice, only including values from slices determined to contain at least 250 intracerebral voxels. Groups deviating by more than 4 standard deviations from the prediction had their data replaced with imputed values. Final interpolation was performed using the Jacobian modulation method to account for signal pile-up/dilution caused by local stretching/compression (Andersson & Sotiropoulos, 2015).

Several confounding time-series were calculated based on the preprocessed DWI: framewise displacement (FD) using the implementation in *Nipype* (following the definitions by Power et al. 2014). The head-motion estimates calculated in the correction step were also placed within the corresponding confounds file. Slice-wise cross correlation was also calculated. The DWI time-series were resampled to ACPC, generating a *preprocessed DWI run in ACPC space* with 2mm isotropic voxels.

### 2.3 Diffusion reconstruction

Reconstruction was performed using *QSIprep* 0.14.3, which is based on *Nipype* 1.6.1 (Gorgolewski et al., 2011; Gorgolewski et al., 2018; RRID:SCR_002502).

#### 2.3.1 DSI Studio reconstruction

Diffusion orientation distribution functions (ODFs) were reconstructed using generalized q-sampling imaging (GQI, Yeh et al., 2010) with a ratio of mean diffusion distance of 1.25. Three tensor values were computed from this fitting that were used in subsequent analyses: axial diffusivity (AD), generalized fractional anisotropy (gFA), and radial diffusivity (RD).

#### 2.3.2 NODDI reconstruction

The NODDI model (H. Zhang et al., 2012) was fitted using the AMICO implementation (Daducci et al., 2015). A value of 0.0017 was used for parallel diffusivity and 0.003 for isotropic diffusivity. This model fit was then used to compute three outcome metrics: orientation dispersion index (ODI), neurite density index (NDI) and free-water fraction (FWF).

#### 2.3.3 MAP-MRI reconstruction

The MAPMRI model (Özarslan et al., 2013) was fitted using the DIPY (Garyfallidis et al., 2014) implementation using default values. Default values of DIPY were used, with the following parameters being set: Radial Order = 6, Laplacian Regularization = True, Laplacian Weighting = 0.2, Positivity Constraint = False, Global Constraints = False, pos grid = 15, pos radius = adaptive, Anisotropic Scaling = True, Eigenvalue Threshold = 0.0001, bval threshold = infinite, dti scale estimation = True, static diffusivity = 0.0007, cvxpy solver = None). Three MAPMRI metrics were used due to their relative independence of correlation mean square diffusivity (MSD), q-space inverse variance (QIV), and return-to-origin probability (RTOP).

### 2.4 Region of interest (ROI) masking

ROI masks were calculated by transforming the Brainnetome (Fan et al., 2016) 246 region atlas from MNI space to subject specific anatomical space through the usage of antsApplyTransforms (Avants et al., 2008) and extracting data for each metric (volume, AD, gFA, RD, ODI, NDI, FWF, MSD, QIV, RTOP) for each ROI. Hippocampal ROI analyses included all four hippocampal ROIs from the atlas (Caudal and Rostral Hippocampus for both hemispheres).

### 2.5 Cognitive testing

Participants included in both Dataset 1 and Dataset 2 were all administered the Rey Auditory Verbal Learning Test (RAVLT) (Rey, 1941). This measure consists of three parts: An initial five presentations of the same 15-word list with the participant being asked to immediately recall as many items as possible from the list, a second immediate recall test after being administered an interference list of 15 different words, and a delayed recall test administered 15 minutes after the second immediate recall test. RAVLT Delay score represents performance on the final part, which is a scale from 0 to 15 based on the number of recalled words. For later analysis, a binarized version of this delay score is used, with individuals with a score greater than 9 are assigned as 1, and those less than or equal to a 9 received a 0.

Participants included in Dataset 2 had several additional metrics available, notably the Trail Making Test and the Digit Symbol Substitution Task. The Trail Making Test Consists of two parts: Part A requires participants to connect circles with numbers in numerical order, while Part B has them connecting alternating numbers and letters in ascending order. The results of the participants’ performance on Part B were used as an outcome metric in the current study. The Digit Symbol Substitution Task (DSST) provides participants with an array of numbers and symbols, and when a number is shown on screen the participant will respond with the associated abstract symbol. This is a timed test, so better performance is indicated by completing more matches in the allotted time. Trails B and DSST are used to assess executive functioning.

### 2.6 Logistic Regression Analysis

For each combination of predictors, a logistic regression analysis was conducted in Scikit-Learn (1.3.0, Pedregosa et al., 2011) with 5-fold cross validation using an L2 penalty function to reduce the risk of overfitting. For analyses focused on the hippocampus, estimates for each metric from each of the four hippocampal ROIs were used, resulting in 4 x the number of metrics for each analysis being predictors in the model (e.g., volume alone results in 4 predictors, Tensors+NODDI+MAPMRI results in 4 x 9 predictors for a total of 36 predictors in the model). For models not predicting binarized age, age was included as an additional controlled variable in each model.

## 3 Results

### 3.1 Hippocampal diffusion metrics easily predict age in older adults

As an initial test of the viability of grey matter diffusion for classification, we median split each dataset by age, replicating the method used in Radhakrishnan et al. (2022). All participants were over age 53, but this splits them into relatively older and younger groups (Dataset 1: 68.38 vs 78.15 mean age; Dataset 2 67.30 vs. 80.94 mean age). When the performed a logistic regression analysis using hippocampal diffusion metrics to classify age group.

Within Dataset 1, 6 sets of metrics were significant predictors of binarized median-split age in older adults: Tensors+NODDI (AUC = .982, p < .001), Tensors+NODDI+MAPMRI (AUC = .982, p<.001), Tensors+MAPMRI (AUC = .857, p < .001), NODDI+MAPMRI (AUC = .821, p<.001), Tensors (AUC = .804, p = .002), NODDI (AUC = .804, p = .002), and MAPMRI (AUC = .714, p = 0.040) (Fig 1A). Similar results were present in Dataset 2, with all 8 sets of metrics being significant predictors of binarized age: Tensors+NODDI (AUC = .855, p<.001), MAPMRI (AUC = .837, p<.001), Tensors+MAPMRI (AUC = .819, p<.001), Tensors (AUC = .804, p<.001), NODDI (AUC = .804, p<.001), NODDI+MAPMRI (AUC = .796, p<.001), Tensors+NODDI+MAPMRI (AUC = .789, p<.001), volume (AUC = .697, p = .026) (Fig. 1B). AUCs appeared to be generally higher in Dataset 1 compared to Dataset 2, potentially due to the relatively lower number of participants relative to predictors resulting in minor overfitting. These results appear to be consistent with previous work that examined these combinations of metrics, which found that Tensors+NODDI combinations provided increased predictive power compared to either metric alone. MAPMRI appears to contribute in a similar way within the first dataset, but the relationship doesn’t appear as present in Dataset 2, with Tensors+NODDI still predicting binarized age with the highest AUC.

**Fig 1.**
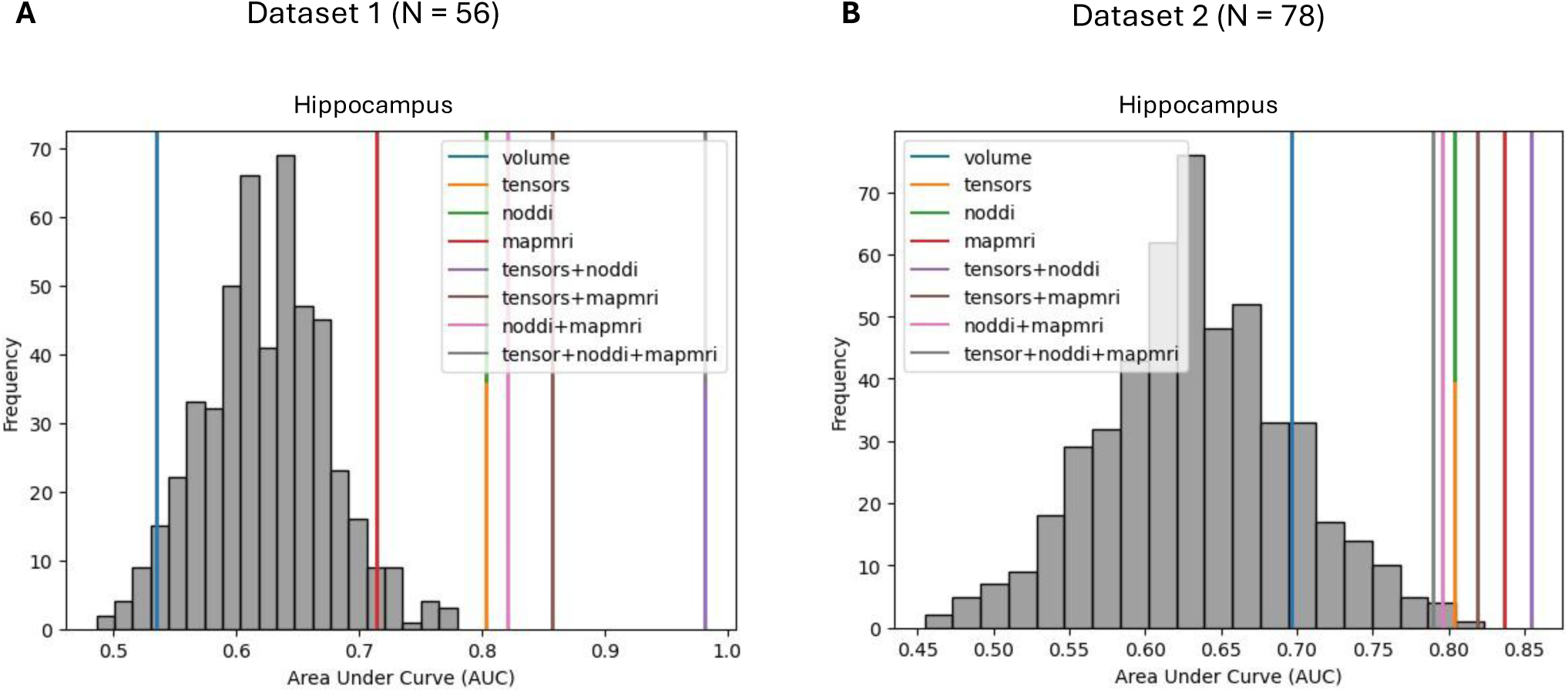
A) Within the first dataset, 6 metrics significantly predicted binarized age via median split with a maximum AUC of .982 for Tensors+NODDI+MAPMRI and Tensors+NODDI and a minimum AUC of .536 for volume. B) Within the ADNI dataset, all metrics significantly predicted age binarized via median split, with a maximum AUC of .855 for Tensors+NODDI and a minimum AUC of .700 for volume. Metric sets with the same AUC are represented by a stacked line.

### 3.2 A combination of hippocampal tensors, NODDI, and MAPMRI metrics strongly predicts RAVLT performance in older adults

We then conducted a similar analysis using hippocampal diffusion metrics to predict binarized RAVLT performance, while controlling for the age of the participants. Within the first dataset, 5 combinations of predictors significantly predict binarized RAVLT performance: Tensors+NODDI+MAPMRI (AUC = 1.0, p<.001), Tensors+MAPMRI (AUC = .925, p<.001), Tensors+NODDI (AUC = .844, p<.001), NODDI+MAPMRI (AUC = .775, p = .012), NODDI (AUC = .744, p = .032) (Fig. 2A). Fewer combinations significantly predict RAVLT performance in Dataset 2, with only 2 combinations being significant predictors: Tensors+NODDI+MAPMRI (AUC = .853, p<.001), Tensors+NODDI (AUC = .790, p<.001) (Fig. 2B). Across both datasets, Tensors+NODDI+MAPMRI is the strongest set of predictors, followed by Tensors+NODDI. This finding of Tensors+NODDI being better than any set of predictors alone is consistent with findings from our previous work (Radhakrishnan et al., 2022). MAPMRI appears to have additional contributions beyond Tensors+NODDI.

**Fig 2.**
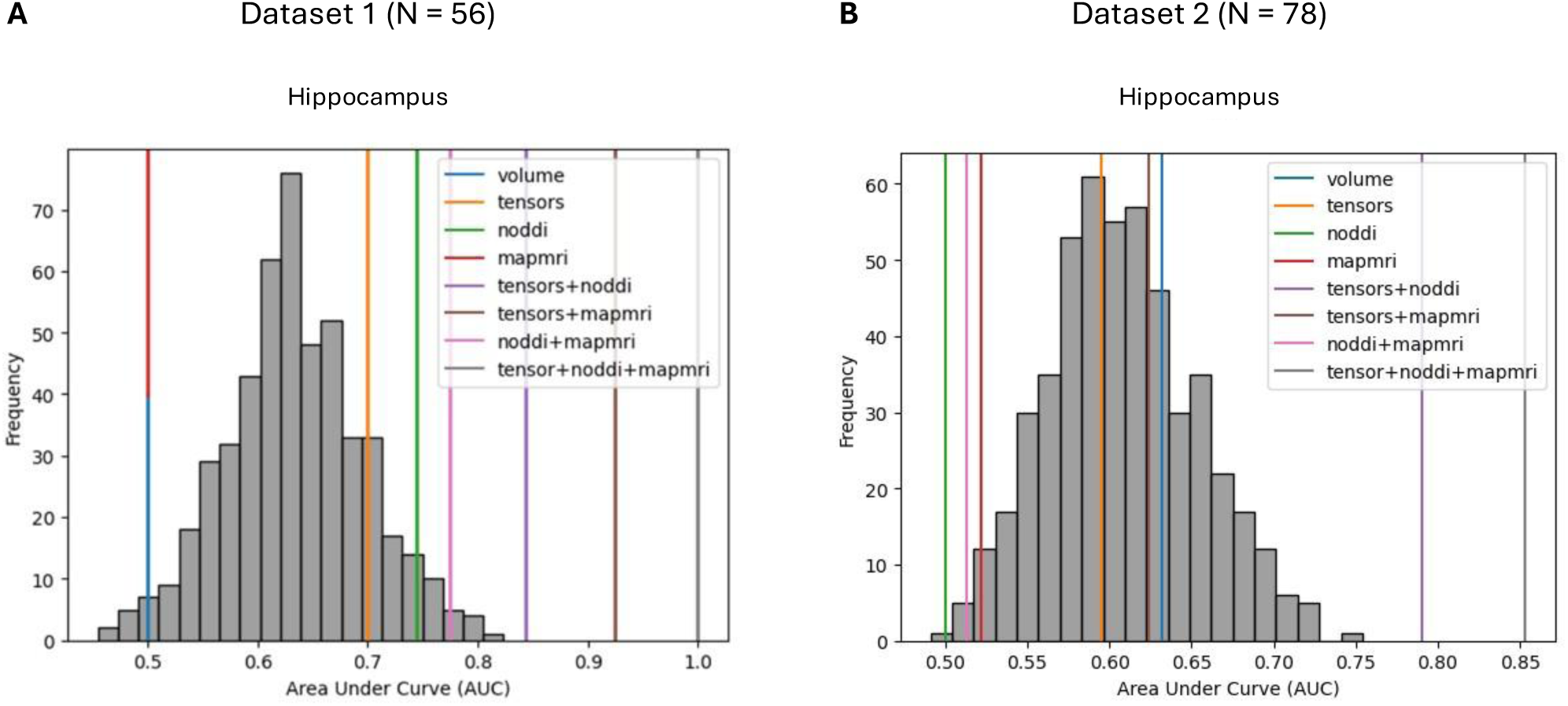
A) Within the first dataset, several combinations of metrics were significant predictors of binarized RAVLT performance (Tensors+NODDI+MAPMRI: AUC=1.000, p<.001; Tensors+MAPMRI: AUC=.925, p<.001, Tensors+NODDI: AUC=.844, p<.001). B) Within Dataset 2, two of the same combinations emerged as significant predictors of binarized RAVLT performance (Tensors+NODDI+MAPMRI: AUC = 0.853, p<.001, Tensors+NODDI: AUC=.790, p<.001). Metric sets with the same AUC are represented by a stacked line.

**Fig 3.**
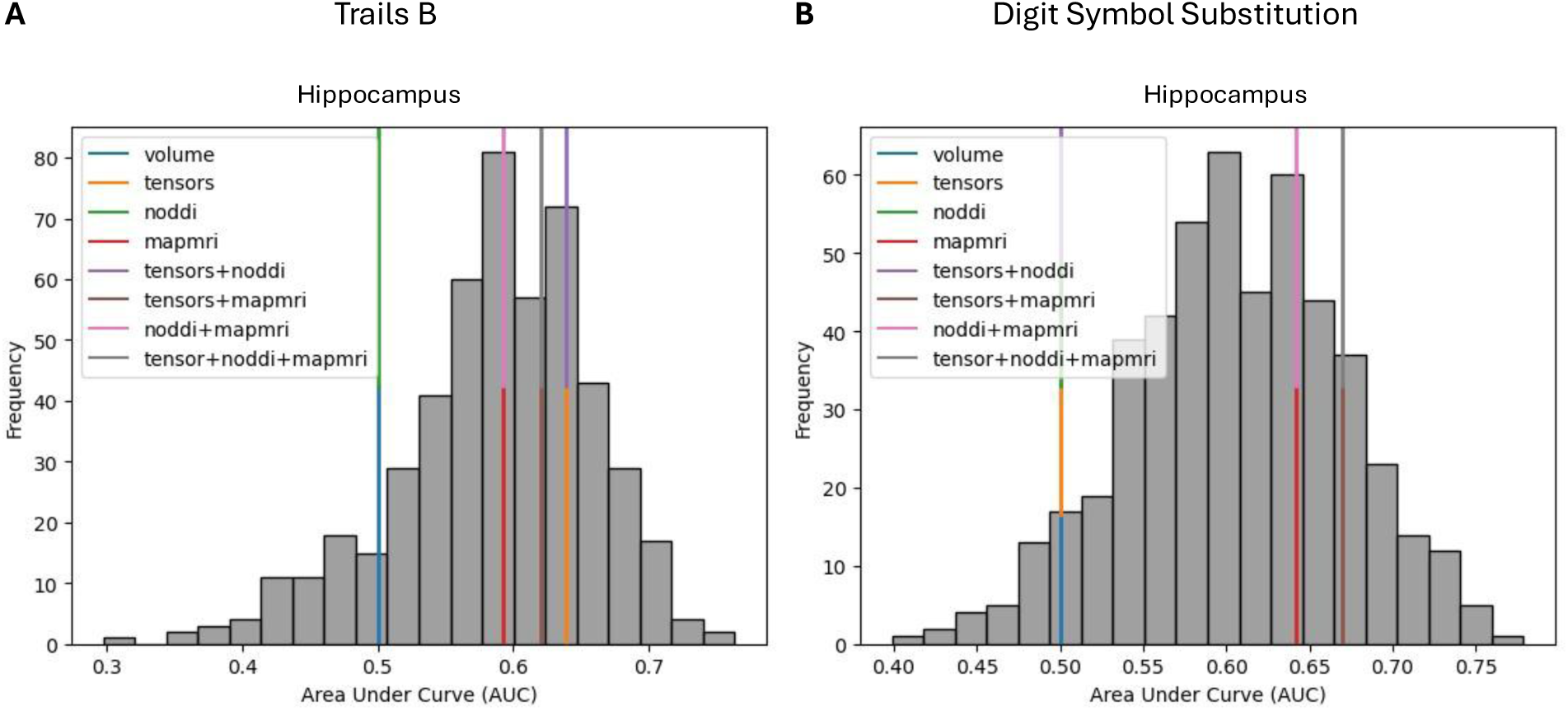
A) Within the Dataset 2, no combinations of diffusion metrics nor volume predicted Trails B performance, with the highest AUC value being for Tensors (AUC = .639, p = .240) and Tensors+NODDI (AUC = .639, p = .240). B) Hippocampal diffusion metrics were also not significant predictors of DSST performance (Tensors+NODDI+MAPMRI: AUC = .669, p = .162; Tensors+MAPMRI: AUC = .669, p = .162). Metric sets with the same AUC are represented by a stacked line.

### 3.3 Hippocampal diffusion appears to uniquely predict explicit memory performance

To examine whether hippocampal diffusion predicted memory performance specific to the RAVLT or just predicted cognition in general, we followed similar procedures to analyze how hippocampal diffusion metrics predicted Trails B memory performance and performance on the Digit Symbol Substitution Task (DSST). For this analysis we focused on Dataset 2, which contained performance metrics for RAVLT, Trails B and DSST for all participants. We found that hippocampal diffusion metrics were not significant predictors of Trails B performance, with the strongest predictors being Tensors (AUC = .639, p = .240) and Tensors+NODDI (AUC = .639, p = .240). Hippocampal diffusion metrics were also not significant predictors of DSST performance (Tensors+NODDI+MAPMRI: AUC = .669, p = .162; Tensors+MAPMRI: AUC = .669, p = .162). These findings indicate that diffusion metrics within the hippocampus are sensitive enough to differentiate between performance on an explicit memory task such as the RAVLT, while not being general predictors of cognition.

### 3.4 Regional diffusion is sensitive to regional differences in task demands

To determine whether the ability of grey matter diffusion to reflect cognition was a more general property, we conducted a search procedure across frontal cortex ROIs present in the Brainnetome atlas (Fan et al., 2016) to identify if any of the ROIs present predicted working memory performance while not predicting memory performance on the RAVLT. One such ROI was identified (Right hemisphere Brodmann Area 9/Brodmann Area 46 ventral portion; represented as A9/46v_R in the Brainnetome Atlas). This region was not a significant predictor of memory performance on the RAVLT, with the strongest predictors being volume (AUC = .567, p = 0.33) (Fig. 4A). Diffusion from this region was, however, a strong predictor of Digit Symbol substitution performance, with the strongest set of predictors being the combination of Tensors+NODDI+MAPMRI (AUC = .631, p = .048) and Tensors+MAPMRI (AUC = .631, p = .048) (Fig. 4B). This indicates that in addition to the sensitivity to specific task demands evidenced in the hippocampus, regional diffusion metrics are also sensitive to different aspects of cognition unique to each region, with at least one region being sensitive to working memory performance but not memory performance in general. Taken together, these findings represent a double dissociation.

**Fig. 4.**
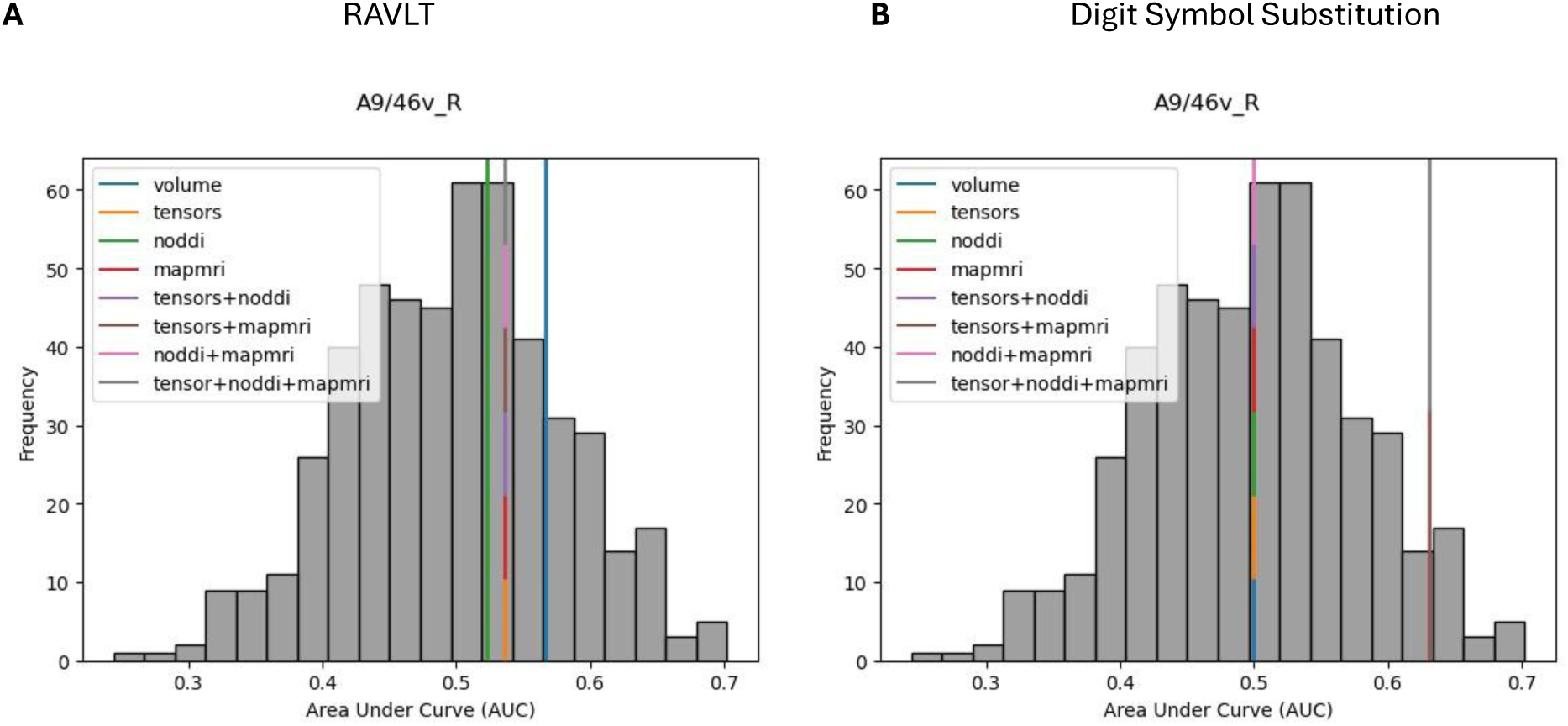
A) Within Dataset 2 we identified an ROI (A9/46v_R) with diffusion metrics that were not able to predict memory performance on the RAVLT (volume: AUC = .567, p = 0.33), B) but was able to predict working memory performance (Tensors+NODDI+MAPMRI: AUC = .631, p = .048; Tensors+MAPMRI: AUC = .631, p = .048). Metric sets with the same AUC are represented by a stacked line.

## 4 Discussion

In the current study, we demonstrated that grey-matter multi-shell diffusion metrics such as NODDI and MAP-MRI can provide significant benefits in predicting age related changes in microstructure in older adults. We reproduced our previous results indicating that a combination of Tensors+NODDI predict age and explicit memory better than either metric alone, generalized these findings across two distinct datasets, and expanded this finding to include MAPMRI and found that combinations of three kinds of diffusion metrics (Tensors+NODDI+MAPMRI) were able to predict both age and memory performance above any combination of two metrics alone.

Consistent with previous work from our lab (Radhakrishnan et al., 2022), we found that all types of diffusion metric (Tensor, NODDI, MAPMRI) from within the hippocampus were strong predictors of age within both of our datasets, and were all substantially better predictors compared to regional volume alone. The finding that even basic information on grey-matter microstructure provided by diffusion imaging being sensitive to age is consistent with previous work by other researchers as well (Leuze et al., 2014; Truong et al., 2014; Aggarwal et al., 2015; Assaf, 2019). Despite the relationships between the types of diffusion metrics, the increase in AUC as more metrics are added is evidence that each of these metrics is conveying at least somewhat distinct information about the hippocampus. Interestingly, despite using a different segmentation of the hippocampus relative to our previous work (our current rostral/caudal segmentation vs specific subfields in Radhakrishnan et al., 2022) we found consistent results with predicting age and memory performance. This might suggest that the level of detail provided by hippocampal segmentation might be lost somewhat when working at lower resolutions with diffusion MRI.

We also were able to provide evidence of a double dissociation within our work. We were able to show that hippocampal diffusion metrics were able to predict memory performance on the RAVLT and that this prediction of cognitive performance didn’t generalize to other cognitive tasks such as Trails B or the DSST. Further, we found that this pattern was differentiable across regions. Several regions were able to predict other cognitive task performance independent of each other, such as Brodmann Area 9/46 being able to predict DSST performance but not performance on other cognitive tasks. This association with Brodmann Area 9 and Digit Symbol performance is well-known (see Silva et al., 2018 for a review). This finding provides even more evidence of the predictive utility of grey-matter diffusion for cognitive and diagnostic purposes.

Our current study has several important limitations. The first limitation is that our data across both samples represents a cross-section of the population. It’s possible that while age is differentiable cross-sectionally within these datasets, that within individuals these changes in metrics over time might not be related in the same ways. As more data is acquired with multi-shell diffusion in datasets such as ADNI with multiple timepoints, this issue can be rectified with longitudinal analysis strategies. Another limitation, as outlined in our previous work, is that we binarized our outcome variables for age and memory performance. While this helps reduce noise and sample bias, we cannot conclusively say whether these same relationships would be observed when predicting absolute age or cognitive performance. A final limitation to note regards the need for multi-shell diffusion data for the advanced diffusion analyses. The need for multi-shell diffusion severely limited the availability of public datasets for use in our study, but with the optimization of multi-shell protocols to be included in future ADNI releases somewhat alleviates this problem.

In conclusion, multi-shell diffusion provides us with a treasure trove of important information on the microstructure of grey matter in the human brain. We have demonstrated that these metrics can be used separately or in tandem to predict both age and cognition in older adults, and that the microstructure of different regions predicts different types of cognitive performance. These metrics provide much value above and beyond volume or tensor-only diffusion and more than justify the extra time required to obtain multi-shell diffusion data.

## 5 Authorship Contribution statement

**Adam Kimbler:** Conceptualization, Methodology, Analysis, Writing – original draft, Visualization. **Craig EL Stark:** Conceptualization, Methodology, Resources, Writing – review and editing, Supervision, Project administration, Funding Acquisition

## 6 Data and Code Availability

Raw and preprocessed data supporting the findings will be made available upon request, made via e-mail to author AK or author CS. The code used for analysis is available here: https://github.com/StarkLabUCI/greymatter_diffusion/

## 8 Acknowledgements

We would like to acknowledge the support of the UCI ADRC and NIH P30 AG066519 award for this project. Data collection and sharing for this project was funded by the Alzheimer’s Disease Neuroimaging Initiative (ADNI) (National Institutes of Health Grant U01 AG024904) and DOD ADNI (Department of Defense award number W81XWH-12-2-0012). ADNI is funded by the National Institute on Aging, the National Institute of Biomedical Imaging and Bioengineering, and through generous contributions from the following: AbbVie, Alzheimer’s Association; Alzheimer’s Drug Discovery Foundation; Araclon Biotech; BioClinica, Inc.; Biogen; Bristol-Myers Squibb Company; CereSpir, Inc.; Cogstate; Eisai Inc.; Elan Pharmaceuticals, Inc.; Eli Lilly and Company; EuroImmun; F. Hoffmann-La Roche Ltd and its affiliated company Genentech, Inc.; Fujirebio; GE Healthcare; IXICO Ltd.; Janssen Alzheimer Immunotherapy Research & Development, LLC.; Johnson & Johnson Pharmaceutical Research & Development LLC.; Lumosity; Lundbeck; Merck & Co., Inc.; Meso Scale Diagnostics, LLC.; NeuroRx Research; Neurotrack Technologies; Novartis Pharmaceuticals Corporation; Pfizer Inc.; Piramal Imaging; Servier; Takeda Pharmaceutical Company; and Transition Therapeutics. The Canadian Institutes of Health Research is providing funds to support ADNI clinical sites in Canada. Private sector contributions are facilitated by the Foundation for the National Institutes of Health (www.fnih.org). The grantee organization is the Northern California Institute for Research and Education, and the study is coordinated by the Alzheimer’s Therapeutic Research Institute at the University of Southern California. ADNI data are disseminated by the Laboratory for Neuro Imaging at the University of Southern California.

